# Combinatorial sequence elements fine-tune mitochondrial protein import to facilitate dual localization

**DOI:** 10.64898/2026.07.31.742112

**Authors:** Youmian Yan, Lianjie Wei, Thomas Schodl, Keri-Lyn Kozul, Maria Behrens, Stephen P. Plassmeyer, Alex S. Holehouse, Natalie M. Niemi

## Abstract

The dual targeting of mitochondrial proteins regulates a host of cellular processes, including metabolism, cofactor biosynthesis, mitophagy, and stress responsiveness. Despite this importance, the mechanisms by which proteins dually localize are incompletely defined. Here, we identify multiple sequence elements that compromise the matrix localization of the phosphatase PPTC7 to facilitate its accumulation at the outer mitochondrial membrane (OMM), where it regulates mitophagy. We find that PPTC7 has a moderately ‘weak’ presequence, but this feature is insufficient to promote dual targeting of a generic cargo protein. Instead, our data suggest that a recently evolved glycine stretch decreases the helical potential of the PPTC7 presequence, weakening its import efficiency in vitro and in cells. Deletion of these glycine residues improves PPTC7 in vitro import and enrichment within the mitochondrial matrix, but only partially suppresses PPTC7-mediated regulation of mitophagy at the OMM. These data suggested additional elements may contribute to PPTC7 dual localization, including its mature phosphatase domain which has robust thermal stability and becomes further stabilized to an import-incompetent state upon binding to its requisite enzymatic co-factor manganese. Simultaneous increases in presequence strength and denaturation of the PPTC7 phosphatase domain are required to promote import in vitro, underscoring the multifactorial challenges associated with its matrix targeting. These data suggest that sequence-specific features can work combinatorially to impart dual-localization capacity to mitochondrial proteins, enabling functions across cellular compartments.

## Introduction

Mitochondria are essential organelles that orchestrate diverse processes such as ATP production, nutrient catabolism, co-factor biosynthesis, and the commitment to cell death^1,2^. Maintaining such an array of functions relies on the coordination of over 1,000 proteins^3,4^, 99% of which are encoded within the nucleus. Thus, intricate mechanisms are required to enable the proper synthesis, targeting, and transport of proteins to and across mitochondrial membranes. The most common mechanism employs an N-terminal cleavable mitochondrial targeting signal, or presequence, to facilitate targeting and membrane translocation^5–7^. However, many proteins utilize less well-understood targeting motifs such as internal or cryptic sequences^8–10^, underscoring the diversity of cellular strategies utilized to achieve organellar protein localization.

Increasingly, there is an appreciation that mitochondria-targeted proteins may not be solely housed at or within the organelle. Multiple examples of dual localization have been described in the literature, with proteins originally thought to be exclusively mitochondrial being found in other subcellular compartments^11–13^, and proteins classified as cytosolic or nuclear harboring a mitochondria-resident fraction^14,15^. This distribution of proteins across subcellular compartments can be dynamic in response to varied stimuli, adding nuance to the sensing and regulation of cellular functions. Indeed, dual-localized mitochondrial proteins have been linked to the regulation of ammonia detoxification^16^, vitamin partitioning^17^, lipid droplet lipolysis^18^, and in genome maintenance^19^. Furthermore, dual-localized proteins have been studied extensively in the context of mitochondrial stress responses, where organellar dysfunction is signaled by proteins whose import is particularly sensitive to alterations in the energetic status of mitochondria^20–23^.

Various mechanisms enable the dual localization of mitochondrial proteins^12,24^. For instance, this could be achieved through the generation of two independent protein products that retain similar functions but differ in their cellular targeting motifs, with processes such as gene duplication, alternative transcription initiation, mRNA splicing, or alternative translation initiation able to generate protein isoforms with or without presequences to achieve multicompartment localization^25,26^. Perhaps a more intriguing scenario involves the dual targeting of a single protein product, which contains sufficient information for mitochondrial targeting but is also capable of partitioning to other cellular locations. The most common mechanism for such dual- localized, single isoform mitochondrial proteins is via a ‘weak’ presequence, which attenuates efficient import into mitochondria to enable the accumulation of a population of the protein at or outside of the organelle. One such dual-localized protein is ATFS-1 in *C*. *elegans*, which translocates to the nucleus to transcriptionally activate the mitochondrial unfolded protein response upon perturbation of its mitochondrial import^20,27,28^. Despite this well-described role of ATFS-1 presequence in stress signaling, it remains unclear how many mitochondrial proteins harbor ‘weak’ presequences, and whether such targeting elements are generally sufficient to promote dual localization. Furthermore, other mechanisms have been identified that may contribute to dual localization, such as protein folding-driven inhibition of mitochondrial import^29,30^, the existence of ambiguous or additional organellar targeting signals within a mitochondrial protein^31^, or recruitment to different compartments via protein binding partners^32^. Despite such a breadth of potential mechanisms suggested to promote dual localization of mitochondrial proteins, many proposed dual-localized proteins have incompletely described modes of localization and regulation. Furthermore, though most models attribute dual targeting to a singular molecular mechanism, multiple sequence features could have additive effects to alter import capacity or protein stability across compartments.

Here, we describe such a multifactorial targeting mechanism for the recently identified dual-localized mitochondrial phosphatase PPTC7^33–35^. Though initially identified as a protein phosphatase in the mitochondrial matrix^36,37^, a second population of PPTC7 was recently discovered at the outer mitochondrial membrane (OMM) to facilitate the turnover of the mitophagy receptors BNIP3 and NIX^33–35,38^. Despite a substantial portion of PPTC7 localizing to both the matrix and OMM in steady state conditions^33–35^, the mechanisms underlying its distribution across compartments are not fully described. Here, we identify multiple sequence elements within PPTC7 that combinatorially diminish its matrix import in vitro and in cells, which likely contribute to its dual localization and thus its ability to dynamically regulate mitochondrial metabolism and mitophagy.

## Results

### PPTC7 has a moderately ‘weak’ presequence that is insufficient to promote dual targeting of GFP

Most mitochondrial proteins harbor N-terminal presequences that facilitate organellar targeting^5,39^. We recently demonstrated that presequences promote variable mitochondrial import efficiency which largely correlates with presequence amphiphilicity^40^. As the PPTC7 presequence has low amphiphilicity relative to three human presequences shown or expected to promote robust import — HSPD1, COX4I1, and COX8A^40–43^, (Figure 1A) — we hypothesized it would drive less efficient import into mitochondria. We leveraged our previously published approach in which variable presequences are fused to a common import cargo, enabling consistent and systematic assessment of presequence strength^40^. Specifically, each presequence was fused to DHFR-HiBiT to generate recombinant fusion proteins that, when incubated with isolated mitochondria containing LgBiT, complement upon matrix import to generate luminescence (i.e., the MitoLuc assay^44–46^) (Figure 1B). This assay allows quantification of various metrics of import efficiency including total imported protein (*b*), maximal translocation rate (*k*), lag time (*t*_1_), and total duration of import (*t*_2_-*t*_1_)^40^ (Figure 1C).

**Figure 1.**
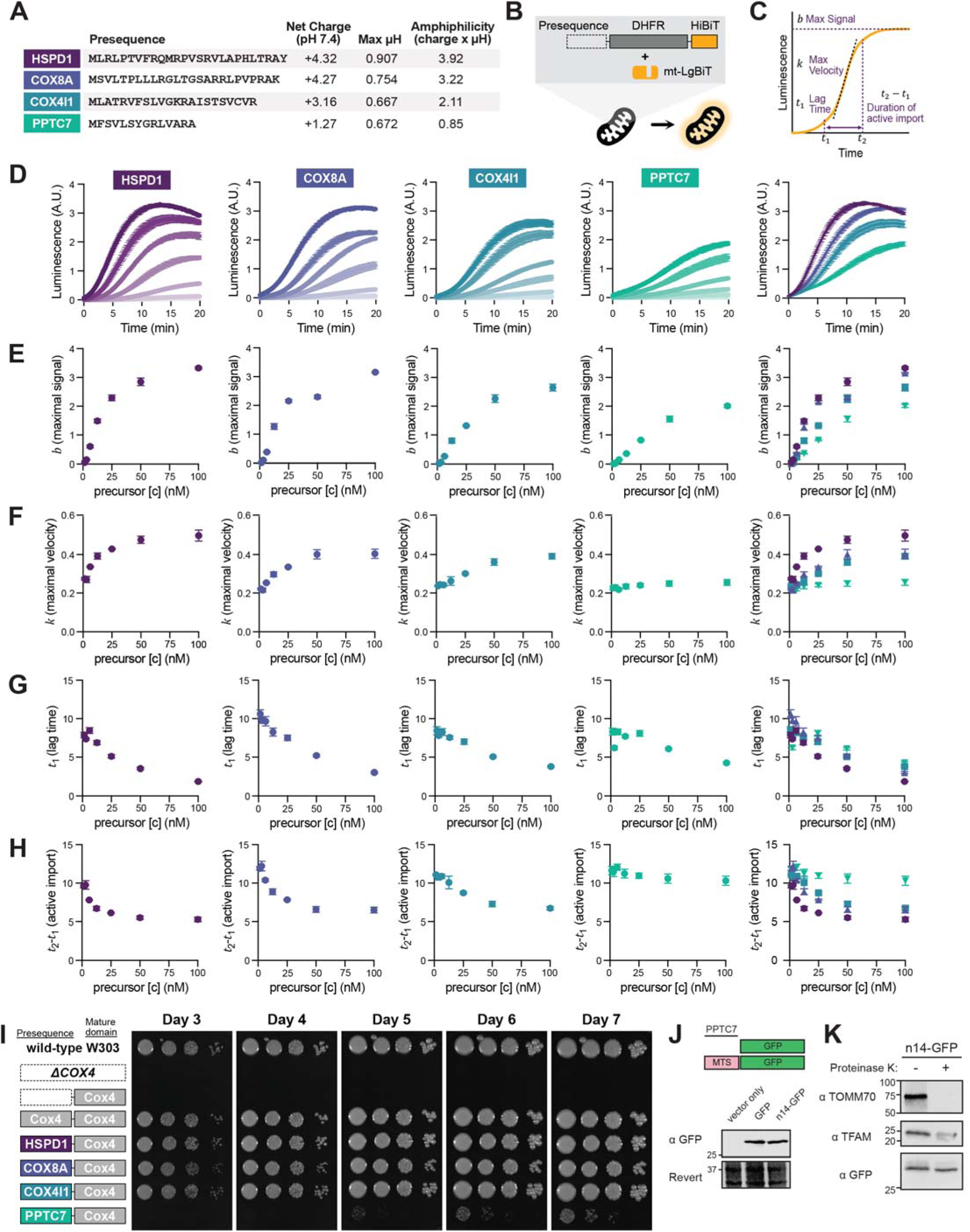
PPTC7 has a moderately ‘weak’ presequence that is insufficient to promote dual targeting of GFP. **(A)** Parameters and sequences of the four human mitochondrial presequences, as defined in reference^39^. **(B-C)** Schematic of MitoLuc (B) and parameters extracted from fitting data to the general logistic function (C). **(D)** MitoLuc measurements of [presequence]-DHFR-HiBiT fusion proteins. Titration curves of precursors at 100 nM, 50 nM, 25 nM, 12.5 nM, 6.25 nM, 3.13 nM, and 1.56 nM; darker shades correspond to higher concentrations. An overlay of import traces of precursors at 100 nM is shown at right. n = 3 independent measurements. Data represented as mean ± standard deviation. **(E-H)** Parameters extracted from the precursor titration (D) by fitting individual curves with the general logistic function. Fitting results shown for total protein imported (E), maximal import rate (F), lag time (G), and duration of active import (H). An overlay of the results is shown at right. n = 3 independent measurements. Data represented as mean ± standard deviation. **(I)** Spotting assay comparing growth of wild-type W303 yeast and Δ*COX4* yeast rescued with vector or the Cox4 mature domain fused to presequences. Serial dilutions of yeast were plated on YPEG (3% ethanol, 3% glycerol) and incubated at 30°C for the indicated timeframe. **(J)** Western blot of GFP or n14-GFP in 293T cells. **(K)** Protease protection assay on isolated mitochondria from 293T cells overexpressing n14-GFP.

As expected, precursors promoted increases in luminescence signal only in mitochondria with an intact membrane potential, indicating bona fide import (Supplemental Figure 1A). Titration of each precursor promoted dose-dependent luminescence traces (Figure 1D) that could be fitted with a general logistic function to extract metrics of import efficiency across concentrations (Supplemental Figure 1B). As expected, the PPTC7 presequence promoted the least efficient import among tested presequences, leading to lower levels of protein imported (Figure 1E), slower rates of import (Figure 1F), extended lag times (Figure 1G), and longer durations of active import (Figure 1H) relative to the three ‘strong’ presequences.

We confirmed these trends in vivo using a Cox4-rescue assay in which presequences, when fused to the mature domain of Cox4, rescue Δ*COX4* yeast respiratory growth proportional to their strength^40,47^. Unexpectedly, our first attempt to express the [PPTC7]-Cox4 fusion failed to promote yeast growth after seven days, and even the ‘strong’ COX8A presequence displayed lagged growth (Supplemental Figure 1C). Upon closer examination, we found these defects may stem from differential codon usage between yeast and humans (Supplemental Figure 1D). We codon-optimized COX8A, which rescued Δ*COX4* respiratory growth similar to the HSPD1 and COX4I1 presequences, while codon-optimized PPTC7 still resulted in a much weaker rescue (Figure 1I). Importantly, each presequence promoted equivalent growth on glucose, indicating growth differences did not stem from loss of viability (Supplemental Figure 1E).

As the PPTC7 presequence promotes relatively inefficient import in vitro and in vivo, we hypothesized it would be sufficient to impart dual localization to a generic cargo protein. We fused the presequence of PPTC7 (i.e., n14) to GFP and found that n14-GFP unexpectedly resolved as a single band when analyzed by Western blotting (Figure 1J). This is inconsistent with the behavior of dual-localized PPTC7, which runs as a doublet of processed and full-length protein^35^. Notably, n14-GFP migrates at the same molecular weight as untagged GFP, consistent with presequence processing and targeting to the mitochondrial matrix. We performed a protease protection assay on isolated mitochondria expressing n14-GFP and found that GFP was resistant to protease treatment, unlike the OMM receptor TOMM70 which was fully digested as expected (Figure 1K). Collectively, these data demonstrate that while PPTC7 possesses a relatively ‘weak’ presequence, this sequence feature is insufficient to impart dual localization of a generic cargo protein, thus suggesting a more complex model underlies the inefficient import of PPTC7 into the mitochondrial matrix.

### An N-terminal disordered region, rather than sensitivity to membrane potential, decreases PPTC7 import efficiency

Multiple proteins achieve dual localization through sensitivity to decreased mitochondrial membrane potential (ΔΨ), including ATFS-1, an effector of the mitochondrial unfolded protein response in *C*. *elegans*^20,27,28^. As the PPTC7 presequence cannot promote dual targeting of GFP in basal conditions, we speculated that it may sense compromised ΔΨ, with its ‘weakness’ exacerbated in conditions of cellular stress^40^. We previously demonstrated that weak presequences, when fused to Cox4, failed to fully rescue the decreased ΔΨ in Δ*COX4* yeast^40^, suggesting that presequences sensitive to ΔΨ may be identified through the Cox4-rescue assay. Consistently, the ATFS-1 and PPTC7 presequences led to similarly poor growth on respiratory media in the Cox4-rescue assay (Figure 2A). We confirmed that the ATFS-1 presequence imparts marked sensitivity to mitochondrial uncoupling in the MitoLuc assay by determining its half maximal inhibitory concentration (IC_50_) of valinomycin, a potassium ionophore that dissipates ΔΨ (Figure 2B). We found that the HSPD1, COX8A, and COX4I1 presequence-containing proteins exhibited valinomycin IC_50_ values roughly five times higher than ATFS-1, consistent with their ‘strong’ import behavior (Figures 2B, C). Unexpectedly, however, the PPTC7 presequence displayed a similar valinomycin IC_50_ as these stronger presequences (Figures 2B, C). To further examine PPTC7 presequence-driven import under perturbed ΔΨ, we used our recently developed PotLuc assay^40^, which couples MitoLuc with a potentiometric dye for simultaneous measurement of protein import and ΔΨ (Figure 2D). These data revealed that the PPTC7 presequence promotes mitochondrial import even in conditions when ΔΨ is compromised (Figure 2E), further supporting its relative insensitivity to diminished ΔΨ. These data indicate that features beyond the presequence of PPTC7 contribute to its dual localization.

**Figure 2.**
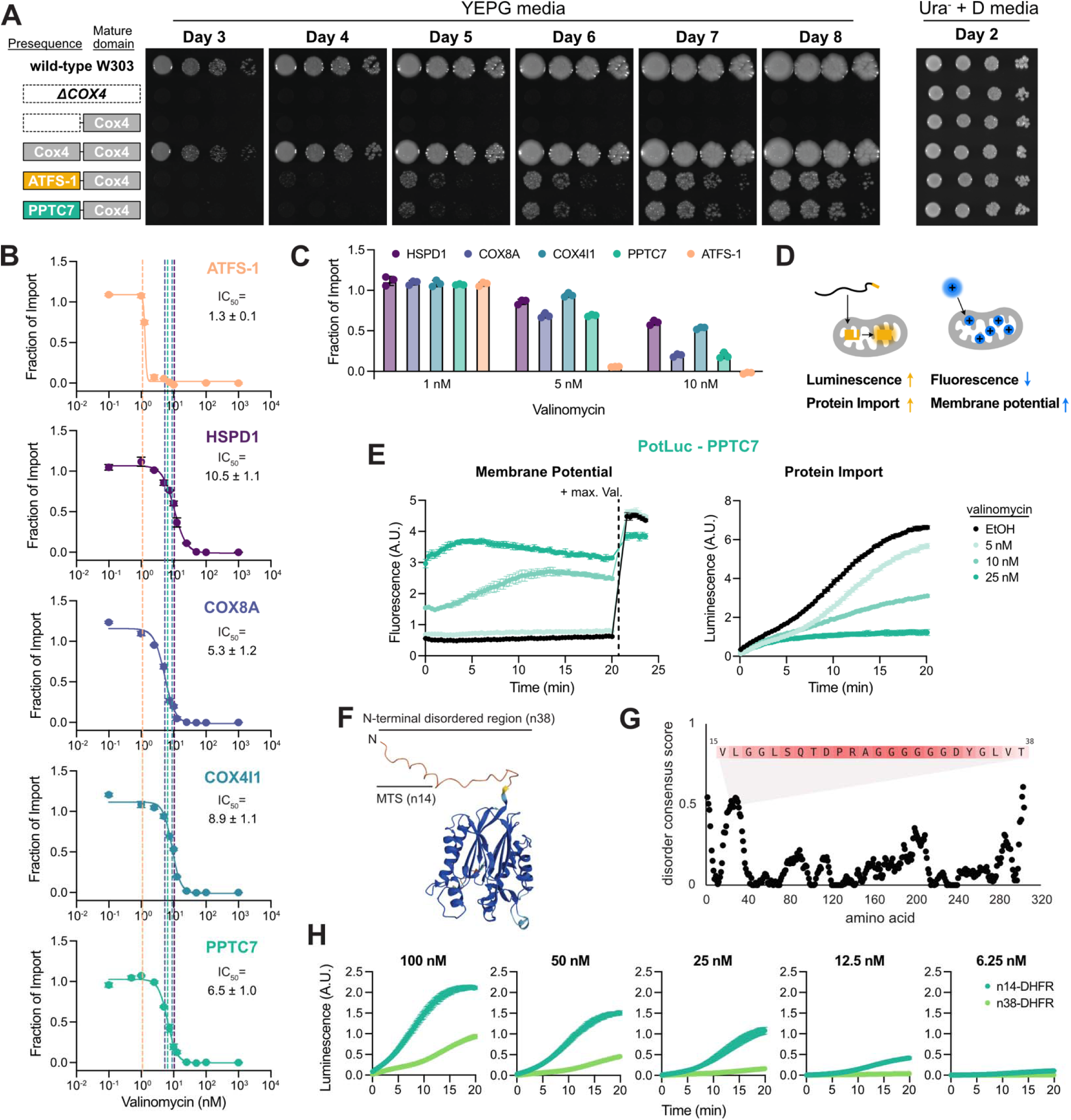
An N-terminal disordered region, rather than hypersensitivity to membrane potential, diminishes PPTC7 import efficiency. **(A)** Spotting assay of wild-type W303 yeast and Δ*COX4* yeast rescued with the Cox4 mature domain fused to [ATFS-1] or [PPTC7] presequence. Serial dilutions of yeast were plated on YPEG (3% ethanol, 3% glycerol) or Ura^-^ (2% glucose) and incubated at 30°C for the indicated timeframe. **(B)** Import of ATFS-1-, HSPD1-, COX8A-, and COX4I1-DHFR-HiBiT in the presence of valinomycin. Fraction of import calculated on maximal signal at each valinomycin concentration relative to vector control in the MitoLuc assay. Data were fitted with a four-parameter logistic function to obtain IC_50_ values. n = 3 independent measurements. Data represented as mean ± standard deviation. **(C)** Comparison of fraction of import at select valinomycin concentrations from (B). **(D)** Schematic of the PotLuc assay. **(E)** Simultaneous measurements of membrane potential and mitochondrial import of PPTC7-DHFR-HiBiT across valinomycin concentrations using PotLuc. n = 3 independent measurements. Data represented as mean ± standard deviation. **(F)** Structure of the full-length PPTC7 predicted by AlphaFold^83^. **(G)** Disorder score for PPTC7 calculated by Metapredict^81^. **(H)** MitoLuc measurements of n14- and n38-DHFR-HiBiT across precursor concentrations. n = 3 independent measurements. Data represented as mean ± standard deviation.

Human PPTC7 is a relatively small protein consisting of a presequence, a disordered linker region, and a PPM-type phosphatase domain (Figures 2F, 2G). This domain architecture is conserved from yeast to humans (Supplemental Figure 2) leading us to predict that the disordered linker might contribute to dual targeting. We tested this by comparing the efficiency of import promoted by the PPTC7 presequence (n14) relative to that driven by the presequence continuous with this disordered stretch (n38) when fused to DHFR. Strikingly, addition of this disordered region markedly reduced import efficiency across precursor concentrations (Figure 2H). Collectively, these data suggest that the PPTC7 presequence is not hypersensitive to altered mitochondrial membrane potential but is instead tuned by a proximal disordered region to diminish its import efficiency.

### The PPTC7 glycine-rich region decreases helical potential and diminishes matrix import in vitro

Examination of the disordered N-terminus of human PPTC7 revealed an unusual stretch of consecutive glycine residues (Figure 3A, Supplemental Figure 3A). Indeed, glycine represents ∼35-40% of the disordered region downstream of the presequence across select mammalian PPTC7 orthologs, leading us to label it the glycine rich region (GRR) (Figure 3B). The unique enrichment and conservation of the glycine residues suggest that the GRR may regulate PPTC7. Presequences tend to form amphiphilic α-helices^47,48^, and glycine is a well-known “helix breaker”, as its small side chain allows for substantial flexibility and free rotation within the peptide bond, disfavoring helical secondary structure^49^. We thus hypothesized that the GRR promotes the acquisition of a heterogeneous ensemble, leading to an entropic penalty that disfavors the propensity of the PPTC7 presequence to adopt a helical conformation. To explore this further, we examined the N-terminal sequence of PPTC7 using SPARROW^50,51^, an analysis package for the quantification and prediction of disordered protein features and properties which reports the likelihood of each residue adopting a helical configuration (Figure 3C). Consistent with our hypothesis, fusing the PPTC7 presequence directly to DHFR (i.e., n14, Figure 3C, teal) increases its predicted helical potential relative to the full, GRR-containing sequence (i.e., n38, Figure 3C, lime green). Furthermore, deletion of the six glycine residues within the GRR (i.e., n38Δ6G) increases presequence helical potential relative to the wild-type n38 sequence (Figure 3C, orange), suggesting this sequence modification would increase protein import capacity. We also used the disordered protein design package GOOSE^52^ to generate three sequence variants, (n38s1, n38s2, and n38s3), which retain identical amino acid composition compared to n38 PPTC7, but alter amino acid placement in the GRR (i.e., between residues 16-38) to increase (n38s1), decrease (n38s2), or maintain (n38s3) presequence helical potential relative to the native N-terminal region (Figure 3C, purple, pink, violet, respectively). We hypothesized that MitoLuc import kinetics would correlate with these predicted helical potentials across PPTC7 presequences.

**Figure 3.**
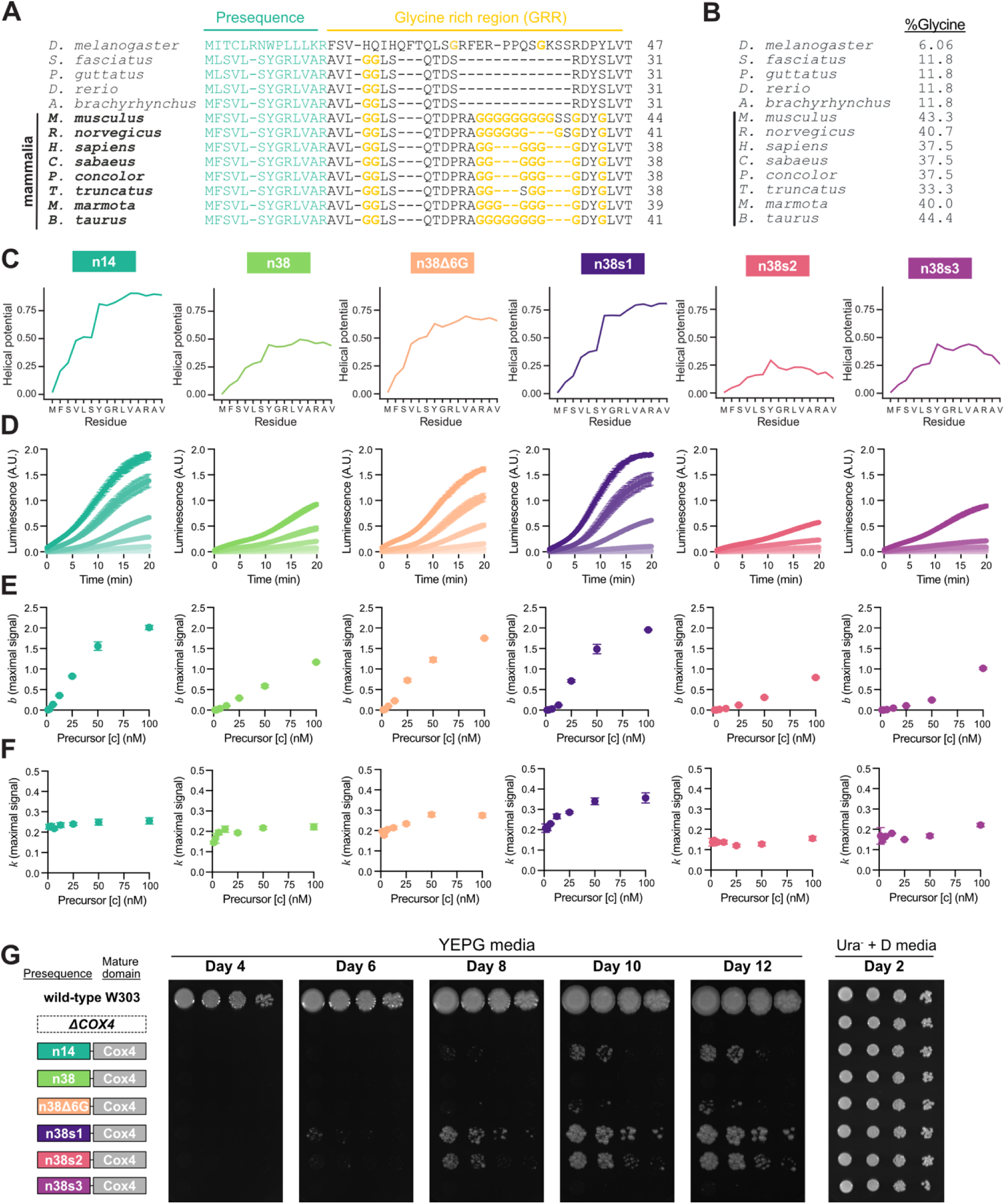
The PPTC7 glycine-rich region decreases import efficiency by tuning presequence helical potential. **(A)** Multiple sequence alignment of the N-terminal region of select PPTC7 homologs. **(B)** Percentage of glycine in the glycine rich region in select PPTC7 homologs. **(C)** Helical potential of the first 15 amino acid residues of the six PPTC7-DHFR-HiBiT fusion proteins calculated by SPARROW^50^. **(D)** MitoLuc measurements of the six [PPTC7]-DHFR-HiBiT fusion proteins across protein concentrations (100 nM, 50 nM, 25 nM, 12.5 nM, 6.25 nM, 3.13 nM, and 1.56 nM), with darker shades corresponding to higher concentrations. n = 3 independent measurements. Data represented as mean ± standard deviation. **(E-F)** Parameters extracted from the precursor titration in (D) by fitting individual curves with the general logistic function. Fitting results for total protein imported (E) and maximal import rate (F) are shown. n = 3 independent measurements. Data represented as mean ± standard deviation. **(G)** Spotting assay of wild-type W303 yeast and Δ*COX4* yeast rescued with Cox4 mature domain fused to PPTC7-derived sequences. Serial dilutions of yeast were plated on YPEG (3% ethanol, 3% glycerol) or Ura^-^ (2% glucose) and incubated at 30°C for the indicated timeframe.

We performed MitoLuc import assays for each of these six constructs across precursor concentrations and fitted the data to a general logistic function (Supplemental Figure 3C). These data revealed that [n38]- DHFR displayed non-saturating, linear trends for total amount of protein imported (*b*), characteristic of weaker presequence behavior in the MitoLuc assay^40^ (Figure 3E). Deletion of the six consecutive glycine residues (i.e., n38Δ6G) largely rescued the import defect seen in [n38]-DHFR (Figure 3D, orange), and the n38Δ6G construct displayed signs of reaching saturation at high precursor concentrations (Figure 3E). We tested the three n38 shuffle variants with synthetically tuned helical potential and found n38s1, predicted to increase presequence helical potential, fully rescued the import defect imparted by the GRR (Figure 3D, purple), giving rise to saturating levels of total protein imported (Figure 3E), higher maximal velocity (Figure 3F), and shorter duration of active import than the native n38 construct (Supplemental Figures 3D, E). The n38s2 variant, which had decreased helical potential, suppressed import to a greater extent than the wild-type GRR (Figures 3D-F, pink). Finally, the n38s3 construct, with similar helical potential to n38, maintained import efficiency relative to the n38 construct at the highest precursor protein concentration, but import capacity declined rapidly at decreased protein concentrations (Figures 3D-F, violet). Similar trends were observed in the Cox4- rescue assay in vivo (Figure 3G), as n14 led to Cox4 rescue but n38 failed to give rise to respiratory growth even after 12 days. Yeast expressing [n38Δ6G]-Cox4 displayed growth, albeit slower than yeast expressing [n14]-Cox4. Furthermore, the stronger shuffle mutant n38s1 promoted more robust growth than n14, and the neutral shuffle mutant n38s3 failed to promote yeast growth, like n38 itself. One inconsistency between the two assays arose with the slower shuffle mutant, n38s2, which has inefficient import in the MitoLuc assay but promoted yeast growth similar to that of n38s1 in vivo. These data suggest that the tuning of helical potential may not be the sole factor affecting presequence strength in vivo, which may be influenced by additional factors such as presequence processing or chaperone association^53^. Nonetheless, these results indicate that the disordered glycine-rich region immediately downstream of the PPTC7 presequence decreases import efficiency and its effects on helical potential contribute to this import inhibition.

### Deletion of the GRR in native PPTC7 disrupts its dual targeting but only partially dampens its inhibition of mitophagy

As the GRR decreases import promoted by the PPTC7 presequence in vitro and in vivo, we hypothesized its deletion would enrich PPTC7 within the mitochondrial matrix. We generated two constructs that lacked either the consecutive glycine residues, Δ6G-PPTC7, or the glycine rich region (i.e., residues 15-32), ΔGRR-PPTC7, and compared their localization to wild-type PPTC7. As expected, overexpressed wild-type PPTC7 ran as a doublet in HeLa FLP-IN cells, consistent with dual targeting (Figure 4A). However, both Δ6G-PPTC7 or ΔGRR-PPTC7 resolve as single bands even in a pseudohypoxic condition that promotes the accumulation of PPTC7 at the OMM^35^ (Figure 4A). These data suggested that disrupting the GRR may diminish its dual targeting due to enhanced matrix import, which we tested via protease protection assays. Consistent with previous findings^33–35^, wild-type PPTC7 was dual-targeted, as the top band of the doublet was digested by externally added proteinase K while the bottom band was protected (Figure 4B). In contrast, deletion of either the consecutive glycine residues or the full GRR resulted in PPTC7 species that were largely protected from proteinase K treatment in isolated mitochondria (Figure 4B), consistent with our data that this region reduces import capacity. These data further suggested that Δ6G-PPTC7 and ΔGRR-PPTC7 may have diminished functionality at the OMM due to their enrichment within the mitochondrial matrix.

**Figure 4.**
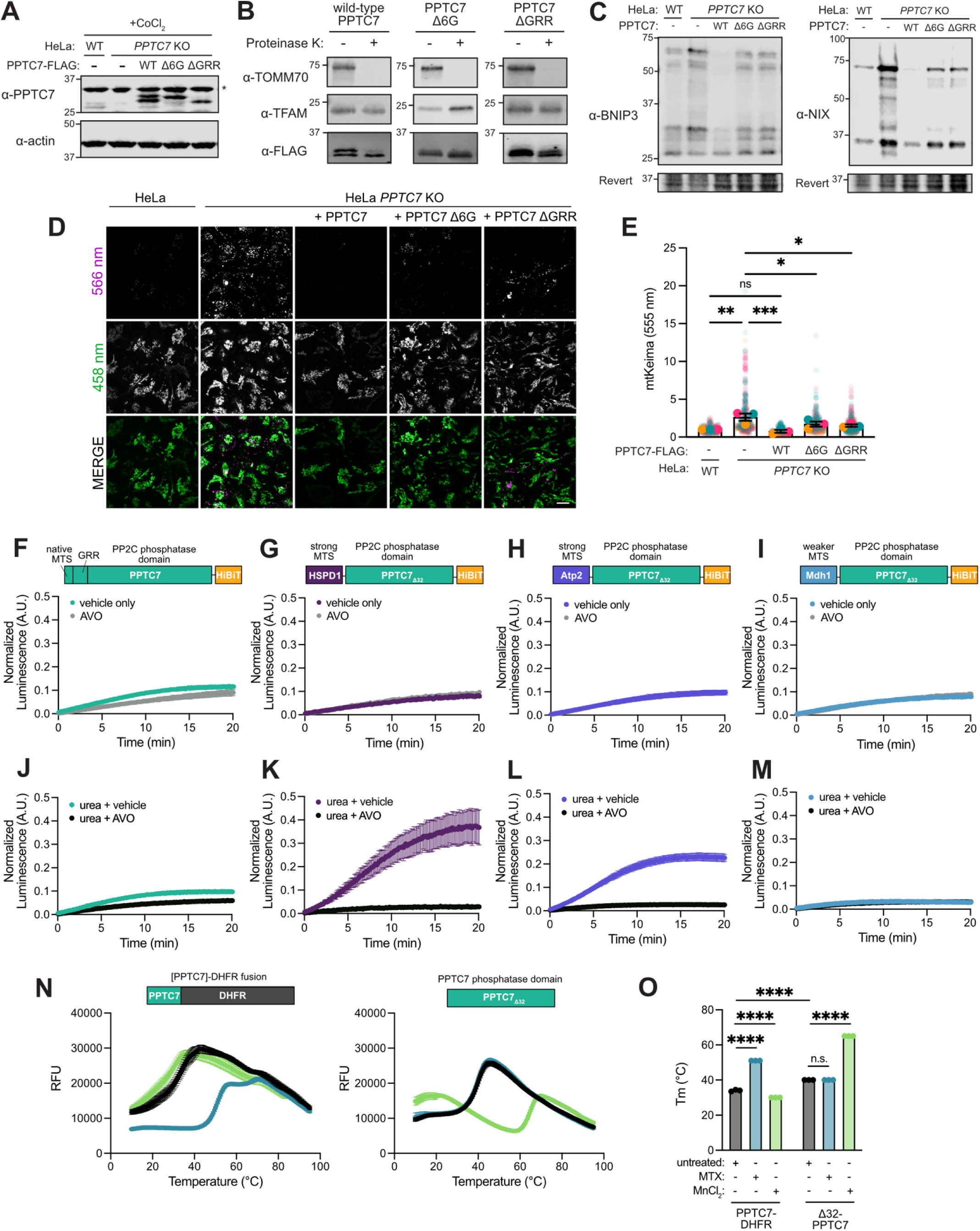
The PPTC7 phosphatase domain is challenging to import. **(A)** Western blot for PPTC7 in wild-type and *PPTC7* KO HeLa FLP-IN TRE-x cells stably expressing wild-type, Δ6G-, or ΔGRR-PPTC7 treated with 200 μM CoCl_2_ for 16 hours. * indicates non-specific band. **(B)** Protease protection assay on mitochondria isolated from 293T cells expressing wild-type, Δ6G-, or ΔGRR-PPTC7. **(C)** Western blot of BNIP3 and NIX in the same cells as in (A). **(D)** mt-Keima analysis of cell lines shown in (A). Acidic mitochondria (top, 566 nm), mitochondria at physiological pH (middle, 458 nm) and merge shown (bottom). Scale bar = 10 μm. **(E)** Quantification of acidic mitochondria as shown in (D). n = 3 independent measurements. Data represented as mean ± standard deviation. n.s., p > 0.05; * p ≤ 0.05; ** p ≤ 0.01; *** p ≤ 0.001; Ordinary one-way ANOVA. **(F-I)** MitoLuc measurements of [PPTC7]-HiBiT (F), [HSPD1]-PPTC7_Δ32_-HiBiT (G), [Atp2]-PPTC7_Δ32_-HiBiT (H), and [Mdh1]-PPTC7_Δ32_-HiBiT (I) into untreated or AVO-treated mitochondria. Luminescence normalized to the maximal signal of the AVO-treated sample. n = 3 independent measurements. Data represented as mean ± standard deviation. **(J-M)** MitoLuc measurements of urea-denatured [PPTC7]-HiBiT (J), [HSPD1]-PPTC7_Δ32_-HiBiT (K), [Atp2]-PPTC7_Δ32_-HiBiT (L), and [Mdh1]-PPTC7_Δ32_-HiBiT (M) into untreated or AVO-treated mitochondria. Luminescence was normalized to the maximal signal of the AVO-treated sample in (F-M) respectively. n = 3 independent measurements. Data represented as mean ± standard deviation. **(O)** Melting curves of [PPTC7]-DHFR and PPTC7_Δ32_ treated with vector, methotrexate (MTX), or MnCl_2_. n = 3 independent measurements. Data represented as mean ± standard deviation. **(P)** Quantification of the melting temperature as shown in (N). n.s., p > 0.05; **** p ≤ 0.0001; Ordinary two-way ANOVA.

As OMM-PPTC7 facilitates the turnover of the mitophagy receptors BNIP3 and NIX, we hypothesized that Δ6G-PPTC7 and ΔGRR-PPTC7 would be less efficient at diminishing the protein levels of these mitophagy receptors than wild-type PPTC7. We tested this by overexpressing each mutant in *PPTC7* KO HeLa FLP-In cells, which have significant elevations in BNIP3 and NIX protein levels (Figure 4C). Consistent with our hypothesis, Δ6G-PPTC7 and ΔGRR-PPTC7 had diminished capacity to decrease BNIP3 and NIX levels relative to wild-type PPTC7 (Figure 4C) and further had compromised ability to suppress levels of mitophagic mitochondria in *PPTC7* KO cells as assayed by the mt-Keima mitophagy reporter (Figures 4D, E). Despite this compromised functionality, both Δ6G-PPTC7 and ΔGRR-PPTC7 maintained partial rescue capabilities of the OMM function of PPTC7. These data suggested that the relatively weak presequence and glycine-rich region of PPTC7 decrease import efficiency in vitro and in cells, but other sequence elements may further contribute to its localization and function at the OMM.

### The PPTC7 phosphatase domain is challenging to import in vitro

We explored whether the mature phosphatase domain of PPTC7 may influence its import and thus facilitate its dual localization. We generated recombinant, full-length PPTC7 containing its presequence, the GRR, and its PP2C phosphatase domain fused to HiBiT at its C terminus and tested its import capacity. Full- length PPTC7 failed to promote detectable ΔΨ-dependent luminescence signal, demonstrating that, when generated recombinantly from *E. coli*, this protein is incompatible with the MitoLuc import assay (Figure 4F). As the PPTC7 presequence and the GRR lead to inefficient import, we generated chimeric constructs that contained the PPTC7 phosphatase domain fused to other presequences, such as the ‘strong’ human HSPD1 and yeast Atp2 targeting motifs, and the yeast Mdh1 presequence, which is similar in strength to the PPTC7 presequence^40^. Interestingly, fusing any of these presequences to the PPTC7 phosphatase domain failed to promote detectable import over an AVO-treated negative control (which disrupts ΔΨ), confirming that this mature domain is challenging to import independent of its N-terminal features (Figures 4G-I).

Protein unfolding has been implicated as rate-limiting in mitochondrial protein import assays in vitro ^54,55^, suggesting we might improve full-length PPTC7 import capacity if we remove this challenge. Unexpectedly, however, urea denaturation of full-length PPTC7 failed to promote import (Figure 4J), though it greatly accelerated the import of [HSPD1]-DHFR as expected (Supplemental Figure 4A). We considered that the N-terminal region of PPTC7 may negatively affect import of PPTC7 even upon unfolding. Indeed, urea treatment of [HSPD1]- and [Atp2]-PPTC7_Δ32_, the two chimeric PPTC7 constructs harboring strong presequences and lacking the GRR, led to observable protein import that could be inhibited by AVO (Figures 4K, L). However, [Mdh1]-PPTC7_Δ32_, which contains a moderately weak presequence but lacks the GRR, displayed no MitoLuc signal over background even after urea denaturation (Figure 4M). These data suggest that PPTC7 is a poor importer in the MitoLuc assay, and both its N-terminal region and its folded domain contribute to this behavior.

We considered that the mature domain of PPTC7 may have high thermal stability, which has a modest inverse correlation with MitoLuc import capacity^40^. Interestingly, PPTC7_Δ32_ had a significantly higher T_m_ than DHFR (Figures 4N, O), consistent with this model. Furthermore, we noted that stabilizing ligands, such as methotrexate (MTX) significantly increase T_m_ of DHFR^56^ (Figures 4N, O). As PP2C phosphatases require divalent cations to function^57–59^, we postulated that enzymatically active PPTC7 may harbor even greater thermal stability upon cofactor binding. We verified that PPTC7_Δ32_ was enzymatically active in a manner dependent on Mn^2+^ (Supplemental Figure 4B), and found that, remarkably, the same concentration of Mn^2+^ that promoted enzyme activity increased the T_m_ of the PPTC7 phosphatase domain by 25°C (Figures 4N, O). Such a stabilizing effect was significantly more pronounced than that of MTX on DHFR (Figures 4N, O), which is well-established to inhibit mitochondrial protein import through stabilization of the DHFR protein fold^56^. Importantly, MTX had no effect on PPTC7 stability, and Mn^2+^ slightly destabilized DHFR, demonstrating ligand specificity (Figures 4N, O). Collectively, our data suggest that multiple, combinatorial sequence elements dictate PPTC7 dual localization. We postulate that together, these properties allow a substantial fraction of PPTC7 to accumulate on the OMM in basal, unstressed conditions to facilitate its role in mitophagic regulation.

## Discussion

There is growing appreciation that many proteins do not reside in a single subcellular compartment. For instance, multiple dual-localized proteins enable mitochondrial stress responsiveness, including PINK1 and DELE1 in mammals^21,22,60–62^, ATFS-1 in worms^20,27,28^, and Mge1 in yeast^23^. These proteins sense mitochondrial dysfunction through alterations in mitochondrial import and/or membrane potential (ΔΨ), which allows a population of each protein to translocate to distinct cellular locations to elicit various responses to mitochondrial stress. Interestingly, proteins not previously linked to stress responses, such as human CblD (gene name *MMADHC*)^17^ and glutamine synthetase (GS) from *G*. *gallus*^16^, also respond to alterations in ΔΨ to facilitate dual localization. CblD accumulates outside of mitochondria upon dissipation of ΔΨ^17^, partitioning cobalamin across cellular compartments. GS from *G*. *gallus* harbors a ‘weak’ presequence that directs this protein to the cytoplasm or mitochondria depending on cell-type differences in ΔΨ^16^. Such examples highlight the utility of ΔΨ-dependent import as a regulatory mechanism to dually target mitochondrial proteins, particularly in conditions of stress.

However, some mitochondrial proteins localize to different compartments in the absence of stress, suggesting alternative molecular mechanisms exist to dually target proteins in basal conditions. The mitochondrial phosphatase PPTC7 is one such protein, as it resides in the mitochondrial matrix^3,36,37^ as well as at the OMM, where it promotes turnover of the mitophagy receptors BNIP3 and NIX^33–35^. This dual localization occurs at steady state across cell lines and tissues, and OMM-localized PPTC7 maintains low levels of BNIP3 and NIX in healthy cells^33–35^, underscoring the need to dually localize PPTC7 when ΔΨ remains intact. Consistently, we demonstrate that the PPTC7 presequence does not have intrinsic hypersensitivity to decreased ΔΨ in vitro, and, although the PPTC7 presequence is moderately ‘weak’ compared to other human presequences, it is insufficient to dually target GFP in cells.

Mammalian PPTC7 homologs evolved a presequence proximal glycine-rich region (GRR) which our data suggest may facilitate its dual localization. Glycine residues disfavor the formation of helices, as their small side chains allow flexibility of the peptide bond. As presequences function as positively charged α- helices, the presence of multiple glycine residues proximal to this region may decrease its helical potential via an entropic penalty, thereby decreasing mitochondrial protein import. Indeed, recent work has implicated the entropic contributions of disordered protein ensembles in regulating other cellular processes, such as membrane curvature sensing^63^, protein interactions^64^, and facilitating protein translocation through bacterial cell walls^65^. Our data suggest that ensemble entropy may additionally tune protein import through membrane translocons. Interestingly, similar findings were reported for aldehyde dehydrogenase (ALDH), as replacement of the native ALDH presequence with the presequence from rhodenase generated a chimeric protein incompatible with import due to a disordered region in ALDH that diminished the helicity of the rhodenase presequence^66^. Our data suggest that PPTC7 exploits a similar mechanism in which the GRR decreases the helical potential of the PPTC7 presequence via an entropic effect. While more work will be required to establish the generality and energetics of this phenomenon, the conserved nature of the PPTC7 domain architecture suggests that this effect is sufficiently important to dictate sequence variation across evolution.

Our data also suggest that the phosphatase domain of PPTC7 likely influences its import capacity, as has been reported in precursors where protein folding within the mature domains can facilitate dual targeting^67^. In yeast, the TCA cycle protein fumarase (Fum1p) partially translocates into the mitochondrial matrix, but folding of its mature domain in the cytosol promotes retrotranslocation and localization in the cytosol^29^. Alternatively, yeast adenylate kinase, Adk1p, largely localizes to the cytoplasm but a minor fraction reaches the mitochondrial intermembrane space. Interestingly, urea-mediated denaturation enriches mitochondrial-localized Adk1p, suggesting protein folding rather than inefficient targeting limits mitochondrial uptake^30^. Like these examples, our data suggest that PP2C phosphatases have a stable fold that is challenging to import. Notably, binding of its requisite enzymatic cofactor Mn^2+^ induced more significant thermal shifts in PPTC7 than MTX-mediated stabilization of DHFR, a classic treatment used to inhibit mitochondrial protein import in vitro and in vivo^56^. These data suggest that, upon binding of metals within the cytosol, PPTC7 likely becomes import incompetent, and this pool of metal-bound phosphatase would be permanently excluded from mitochondrial matrix. Such a mechanism may obligate a population of PPTC7 to the OMM to facilitate the constitutive turnover of BNIP3 and NIX.

Metal binding is a conserved feature of PP2C phosphatases, and multiple PP2C phosphatases localize to mitochondria from yeast to mammals^68^. Interestingly, at least four mitochondrial PP2C phosphatases (including PPTC7) have reported dual localization. The BCKDH phosphatase PPM1K localizes not only to the mitochondrial matrix but also the cytosol to dephosphorylate ACLY and promote hepatic lipogenesis^69^. In yeast, Ptc5p dually targets to mitochondria and peroxisomes via a ‘tug-of-war’ mechanism between the import complexes of each organelle^70^. Finally, Ptc7p, the yeast homolog of PPTC7, displays dual localization but via alternative splicing^25^. Full-length, unspliced Ptc7p localizes to endoplasmic reticulum^71^ or nuclear envelope^25^ to perform unknown functions, whereas spliced Ptc7p localizes to the mitochondrial matrix^25^ to dephosphorylate proteins involved in the TCA cycle^72^, protein import^37^, and CoQ biosynthesis^73^. Notably, both Ptc5p and Ptc7p may overcome challenges in protein unfolding or cofactor- mediated stabilization via co-translational import^71^. However, it remains unclear how dual-localized proteins such as PPTC7 would accumulate outside of mitochondria if fully engaged in co-translational import, suggesting alternative mechanisms likely contribute to their distribution across cellular compartments.

Collectively, our work suggests that human PPTC7 harbors unique, combinatorial sequence features that disallow its efficient targeting into mitochondrial matrix. Importantly, such sequence features would allow the constitutive accumulation of PPTC7 at the OMM even in basal conditions, representing a distinct targeting mechanism in contrast to those that are well characterized in mitochondrial stress responsiveness. Further work will be required to understand the extent to which ‘weak’ presequences, altered presequence helicity, and robust thermal stability of mature protein folds contribute to variability in import efficiencies of the hundreds of mitochondrial proteins expressed across cell types and organisms. Such insights may reveal additional layers of complexity in the regulation of mitochondrial homeostasis and intercompartmental crosstalk.

## Acknowledgements

This work was supported by the National Institutes of Health (R35GM151130 to N.M. Niemi), the National Science Foundation (NSF CAREER award #2338129 to A.S.H.), and start-up funds from the Department of Biochemistry & Molecular Biophysics (to N.M. Niemi). K-L.K was supported by a Postdoctoral Fellowship from the United Mitochondrial Disease Foundation via generous support from the Kamaria Satcher Fund for Kearn’s Sayer Syndrome. T.S. was supported by an Olin Fellowship from Washington University in St. Louis. We thank members of the Niemi laboratory as well as Jonathan Friedman (UTSW) for helpful discussions and feedback on this work. We thank the Robertson lab (WUSM) for sharing their sonicator for experiments associated with this work. Experiments were performed using a Zeiss LSM980 AiryScan 2 in part through the use of Washington University Center for Cellular Imaging (WUCCI) supported by Washington University School of Medicine, The Children’s Discovery Institute of Washington University and St. Louis Children’s Hospital (CDI-CORE-2015-505 and CDI-CORE-2019-813) and the Foundation for Barnes-Jewish Hospital (3770 and 4642). The Zeiss LSM 980 Airyscan Confocal Microscope which was purchased with support from NIMH grant S10MH126964.

## Author Contributions

Y.Y., L.W., and N.M.N. conceived the overall project and its design. Y.Y., L.W., K-L.K., T.S., M.B., S.P.P., A.S.H. and N.M.N. generated and validated tools, performed experiments, performed formal analyses, curated data, and/or assisted with data analysis and interpretation of data. Y.Y., L.W., and N.M.N. wrote the manuscript and generated the associated figures. All authors reviewed and edited the manuscript.

## Methods

### Molecular Cloning

For purification of recombinant proteins, the backbone vectors used were obtained from the laboratory of David Chan^74^ (pET28-Mff(1-61)-PP-GST; #73042; Addgene) and Stephen Fuhs and Tony Hunter^75^ (pGEX6P-1-NME2 H118Y). For protein expression in yeast, the backbone vector used was obtained from the laboratory of David Pagliarini^72^ (pJR13019). Standard cloning procedures with PCR, restriction enzyme digestion, ligation, and bacterial transformation (catalog #2988J; New England Biolabs) were performed to create the desired plasmids. For codon optimization, the online tool developed by Integrated DNA Technologies was used. For stable transfections in HeLa FLP-IN TREx system as well as transient transfections in HEK293T WT cells, pcDNA5 FRT/TO plasmids were generated using Gateway cloning. Briefly, constructs were PCR amplified with primers containing attB1 or attB2 sequences or ordered as gBlocks from Integrated DNA Technologies. These fragments were incubated with pDONR221 (a gift from Julia Pagan) and BP clonase for recombination. Positive constructs were incubated with LR clonase and the pcDNA5/FRT/TO-Venus-Flag-Gateway destination vector, which was a kind gift from Jonathon Pines^76^ (#40999; Addgene). All cloned plasmids were validated via Sanger sequencing.

### Yeast cultures

The WT haploid W303 (his3 leu2 lys2 met15 trp1 ura3) *Saccharomyces cerevisiae* strain was a gift from the laboratory of David Pagliarini. The generation of Δ*COX4* yeast strain and the procedure for yeast transformation and spotting assay have been described^40^. YPEG (1% yeast extract, 2% peptone, 3% ethanol, and 3% glycerol) or synthetic minus uracil (catalog #D9535; US Biologicals) with 2% glucose plates were used for spotting assay. For each yeast strain, 10^4^, 10^3^, 10^2^, and 10^1^ cells were plated and incubated at 30°C for the indicated length of time. The ChemiDoc MP Imaging System (Bio-Rad) and Image Lab Touch Software (Bio-Rad, version 3.0.1.14) were used for imaging the yeast plates.

### Purification of recombinant proteins

Detailed procedures have been previously published^40,46^. Briefly, BL21 (DE3) competent *E. coli* cells (catalog #C2527I; New England Biolabs) were transformed with the target plasmid. The cells were grown in LB media at 37°C with shaking (225 rpm). Protein expression was induced by addition of 1 mM IPTG when the bacterial cultures reached mid-log phase, and the cultures were transferred to 18°C to grow for another 18 hours. Cells were pelleted by centrifugation and then resuspended in GST lysis buffer (1x PBS, 1% [v/v] Triton X-100, 5% [v/v] glycerol, 1 mM DTT) on ice. Sonication was performed to lyse the cells, and the soluble fraction was collected after centrifugation at 14,000 *g* for 30 min at 4°C. Glutathione resin (catalog #NC1057345, Thermo Fisher Scientific) was washed three times with 10x bed volume of ice-cold GST lysis buffer and then incubated with the supernatant collected for 2.5 hours at 4°C. The resin was washed with 10x bed volume of GST lysis buffer three times and then 10x bed volume of PreScission Protease buffer (20 mM Tris-HCl, 150 mM NaCl, 0.5 mM EDTA, and 1 mM DTT, pH 7.4) three times at 4°C. The pelleted resin was resuspended in 1x bed volume of PreScission Protease buffer with the PreScission Protease. After overnight incubation at 4°C, the supernatant was collected. Centrifugal concentrators with a 10K molecular weight cutoff (catalog #VS2001; Vivaproducts) was used to concentrate the proteins. Protein aliquots were stored at -80°C until usage. Protein samples and BSA standards (catalog #PI23225; Thermo Fisher Scientific) were analyzed by SDS-PAGE to calculate the concentrations. For the three chimeric [presequence]-PPTC7 constructs, the concentrations were too low to be accurately approximated, but their expression was confirmed through Western blotting.

### Isolation of crude mitochondria from yeast

Detailed procedures have been previously described^40,46^. Yeast cells were grown in synthetic minus uracil media containing 2% glucose at 30°C overnight with shaking (225 rpm). The overnight cultures were then diluted in YPG media (1% yeast extract, 2% peptone, and 3% glycerol) to reach a concentration of 8 x 10^5^ cells/mL. After the culture reached mid-log phase, 1% galactose was added to induce the expression of mt- LgBiT for 3 hours. After collecting the cells through centrifugation, the pellet was resuspended in DTT buffer (100 mM Tris-H_2_SO_4_, 10 mM DTT, pH 9.4) and incubated at 30°C for 20 min with shaking (80 rpm). The cells were pelleted and then washed with zymolyase buffer (1.2 M sorbitol, 20 mM potassium phosphate, pH 7.4). After pelleting the cells again, the cells were incubated in zymolyase buffer containing zymolyase (catalog #NC0497252; Thermo Fisher Scientific) for 45 min at 30°C with shaking (80 rpm). After another wash with zymolyase buffer, the cells were resuspended in ice-cold homogenization buffer (0.6 M sorbitol, 10 mM Tris- HCl, 1 mM EGTA, 1 mM PMSF, 0.2% [w/v] fatty acid-free BSA, pH 7.4). A glass Dounce homogenizer was used to lyse the yeast cells on ice. The lysates were centrifuged at 1,500 *g* and then 4,000 *g* for 5 min at 4°C to pellet cell debris and the nucleus. The supernatant was then centrifuged at 12,000 *g* for 15 min at 4°C to pellet the mitochondria, which was then resuspended in SEM buffer (250 mM sucrose, 1 mM EGTA, 10 mM MOPS-KOH, pH 7.2) on ice with wide-bore tips. The Pierce 660 nm Protein Assay Kit (catalog #22662; Thermo Fisher Scientific) was used to measure the mitochondrial protein concentration. Aliquots were flash- frozen in liquid nitrogen and stored at -80°C until usage.

### Cell Culture

HEK293T cells were acquired from the American Type Culture Collection (Manassas). HeLa FLP-IN TREx cells stably expressing mt-Keima were a kind gift from Dr. Julia Pagan. *PPTC7* KO HEK293T cells and *PPTC7* KO HeLa mtKeima cells were generated using CRISPR-Cas9 technology as previously described^38,77^. Cells were cultured in growth media (Delbecco’s Modified Eagle Media supplemented with 10% heat inactivated Fetal Bovine Serum and 1x penicillin/streptomycin). Cells were grown in a temperature- controlled incubator at 37°C and 5% CO_2_. Transient plasmid transfection into HEK293T wild-type and *PPTC7* KO cells was performed with polyethylenimine (PEI) for 24 to 48 hours. Stable plasmid transfection into HeLa FLP-IN TREx cells were performed in the presence of pOG44 at a ratio of 0.5 μg pcDNA5 to 2 μg pOGG44. Transfections were performed with FuGENE 6 (catalog #E2691; Promega) per manufacturer’s directions for 24-48 hours before selection with 400 μg/mL hygromycin B (Thermo Fisher Scientific) for approximately 7–10 days. To induce expression of wild-type or mutant PPTC7, cells were treated with 1 μg/mL doxycycline for 24-32 hours. To stimulate pseudohypoxic conditions, cells were treated with 200 μM CoCl_2_ for 16 hours prior to harvesting.

### Protease protection assay on isolated mitochondria

A previously published protocol was followed^78^. Briefly, HEK293T WT cells were seeded in 15-cm plates. On day 2, the plasmids carrying different PPTC7 constructs were transfected with PEI. After 24 hours, cells were then harvested by scraping in PBS and spun down at 600 *g* for 10 minutes at 4°C. To obtain a mitochondrial- enriched fraction, cells were resuspended in mitochondrial isolation buffer (0.1 M Tris-MOPS, pH 7.4, 0.1 M EGTA, 1 M sucrose in water) and homogenized at 1000 rpm with a Potter-Elvehjem tissue homogenizer at 4°C. The resulting mixture was then centrifuged at 600 *g* for 10 minutes at 4°C, and the supernatant was then kept and recentrifuged at 7,000 *g* for 10 minutes at 4°C. The resulting pellet was the washed and resuspended with mitochondrial isolation buffer and quantified by using the Pierce 660nm Protein Assay Kit (catalog #22662; Thermo Fisher Scientific). After crude mitochondrial isolation, the mitochondria-enriched fraction was diluted in mitochondrial isolation buffer (20 mM HEPES pH 7.5) before incubation on ice for 15 minutes. Afterwards, proteinase K (50-100μg/mL final concentration; GoldBio) was added as necessary per reaction. The reaction mixtures were incubated on ice for 10 minutes before quenched by the addition of 5 μM final concentration of phenylmethylsulfonyl fluoride (PMSF). The reaction mixture was chilled on ice for another 5 minutes before centrifugation at 10,400 *g* for 15 minutes at 4°C. 12.5% [w/v] final concentration of trichloroacetic acid was added to the reaction mixtures, and these reaction mixtures were heated at 65°C for 10 minutes before incubation on ice for 45 minutes. The mixtures were then undergone centrifugation at 21,100 *g* for 5 minutes, and the supernatant was discarded. The resulting pellets were then washed twice with ice-cold acetone with centrifugation at 21,100 *g* for 5 minutes in between. The washed pellets were then allowed to dry completely before resuspension in 1x sample buffer in RIPA and were subjected to SDS- PAGE and subsequent immunoblotting analysis.

### Western blotting

Cells were lysed with radioimmunoprecipitation buffer (RIPA; 0.5% w/v sodium deoxycholate, 150 mM sodium chloride, 1.0% v/v IGEPAL CA-630, 1.0% sodium dodecyl sulfate (SDS), 50 mM Tris pH 8.0. 1mM EDTA pH 8.0 in water) supplemented with 1x protease inhibitor cocktail (0.5 μg/mL pepstatin A, chymostatin, antipain, leupeptin, and aprotinin) and 1x phosphatase inhibitor cocktail (0.5 mM imidazole, 0.25 mM sodium fluoride, 0.3 mM sodium molybdate, 0.25 mM sodium orthovanadate, and 1 mM sodium tartrate) unless otherwise specified. After generating cell lysates, samples were clarified by centrifugation (21,100 x g) at 4°C for 10 minutes, snap frozen in liquid nitrogen, and stored at -80°C before use. All samples were quantified with the bicinchoninic acid (BCA) assay kit (Thermo Scientific). Lysates were mixed with 5x sample buffer (312 mM Tris-Base, 25% w/v sucrose, 5% w/v SDS, 0.05% w/v bromophenol blue, 5% v/v β- mercaptoethanol, pH 6.8) and boiled at 95°C for 10 minutes. Lysates (20-40 μg) were run on SDS-PAGE gels with Precision All-Blue Protein Standards (catalog #1610373; Bio-Rad) before being transferred onto nitrocellulose membranes. Total protein were stained by Revert 700 Total Protein Stain (catalog #926-11021; LI-COR Biosciences). After blocking with 2% BSA or 3% nonfat dairy milk in TBS-T, membranes were incubated with following primary antibodies diluted 1:1000 in 2% BSA or 3% nonfat dairy milk in TBS-T at 4°C: GFP (overnight; catalog #2555; Cell Signaling Technology), TFAM (overnight; catalog #44060; Cell Signaling Technology), TOMM70 (overnight; catalog #65619; Cell Signaling Technology), PPTC7 (48 hour incubation; catalog #NBP190654; Novus), BNIP3 (48 hour incubation; catalog #44060; Cell Signaling Technology), NIX (48 hour incubation; catalog #12396; Cell Signaling Technology), β-actin (overnight; catalog #3700; Cell Signaling Technology). Membranes were washed 2-3x with TBS-T for 5 minutes per wash and incubated with corresponding fluorophore-conjugated antibodies for 30 minutes at room temperature. Anti-680 or anti-800 conjugated mouse or rabbit antibodies (catalog #926-32210 and 926- 32211; LI-COR Biosciences) were used for detection. Membranes were then washed 2-3x with TBS-T for 5 minutes per wash and were imaged with a LiCOR OdysseyFC instrument using Image Studio software (LiCOR; version 5.2).

### Microscopy for mt-Keima analysis

Quantification of mitophagy levels using the mt-Keima assay was performed following previously published protocols^38,79,80^. Live cells were imaged by excitation at 458 nm (mitochondrial signal) and 561 nm (mito- lysosomal signal), and emission at 620 nm on a Zeiss LSM980 AiryScan 2 microscope with a 63x high NA oil immersion objective. The environmental chamber was set to 37 °C with 5% CO2. The tiling position feature of randomized positions in the field of vision was used to obtain images. The mito-lysosomal signal at 561 nm was quantified using ImageJ2 software. Single cells were defined into distinct regions of interest (ROIs), which were then split into different channels for threshold processing and measurement using ImageJ2. The results were normalized to the wild-type control, and graphed using Graphpad Prism software (v.10.6.1).

### MitoLuc assay

Detailed protocols have been published previously^40,46^. Briefly, 2x import buffer (500 mM sucrose, 160 mM KCl, 2 mM potassium phosphate, 10 mM MgCl_2_, 20 mM MOPS-KOH, 0.2% [v/v] Prionex reagent, pH 7.2) was prepared on ice. Mixture 1 (62.5 μg/mL of mitochondria in 1x import buffer) and mixture 2 (5x precursor proteins and 1.25x Nano-Glo luciferase assay substrate (catalog #N2012; Promega) in 1x import buffer) were prepared separately on ice. For AVO treatment, 12.5 μM antimycin A (catalog #A8674; MilliporeSigma), 1.25 μM valinomycin (catalog #V0627; MilliporeSigma), and 25 μM oligomycin (catalog #O4876; MilliporeSigma) were added to mixture 1. For valinomycin titration, valinomycin was added to mixture 1 at 1.25x of target concentration. For urea denaturation, urea was added to mixture 2 to reach a final concentration of 6 M. To start the import reaction, 100 μL of mixture 1 and 25 μL of mixture 2 were mixed on a white flat-bottom 96- well plate (catalog #30196; Pro Lab Supply). The BioTek Synergy LX Multimode Reader (Agilent Technologies) was used to measure the luminescence signals with its luminescence filter, and BioTek Gen5 Software (Agilent Technologies, version 3.11) was used for data acquisition. The program was set to measure luminescence (acquisition time = 0.2 sec, gain = 100) every 8 sec with linear shaking of the plate for 1 sec in between. For data analysis, measurements after reaching the maximal luminescence signal for each well were excluded, and the import curves were fitted with the general logistic function:

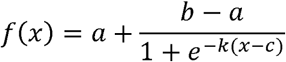

Lag time and duration of active import were calculated by solving for the points where the third derivative of the logistic function was equal to 0. Model fitting through nonlinear regression and derivative calculation were performed using MATLAB (MathWorks, version R2025b). For the valinomycin titration in Figure 2, the import of 100 nM of ATFS-1-DHFR, 25 nM of HSPD1-DHFR, 50 nM of COX8A-DHFR, 50 nM of COX4I1-DHFR, or 100 nM PPTC7-DHFR was quantified at varied valinomycin concentrations. The fraction of import was calculated using the equation

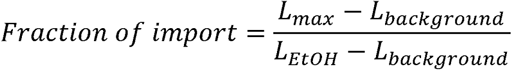

Where L_max_ was the maximal luminescence signal at each condition, L_background_ was the maximal luminescence signal at 1 μM valinomycin, and L_EtOH_ was the maximal luminescence signal from the EtOH-treated trace. The results were fitted with the “[Inhibitor] vs response - Variable” model in Prism (GraphPad, version 11.0.2) to obtain the IC_50_ values, and standard deviations were calculated from the 95% confidence interval.

### PotLuc assay

A detailed procedure has been published previously^40^. Briefly, the procedure was the same as the MitoLuc assay described above with the following deviations. Mixture 1 contains 1.25 μM of DiSC_3_(5) and valinomycin that was 1.25x of the target concentration. A clear bottom 96-well plate was used for loading the samples. The BioTek Cytation 5 Cell Imaging Multimode Reader (Agilent Technologies) and the BioTek Gen5 software (Agilent Technologies, version 3.15) were used for collecting the data. The program was set to measure luminescence and fluorescence (excitation at 622 nm, emission at 670 nm) every 20 sec with 1 sec of linear shaking of the plate in between. After 20 min of measurement, 1 μM of valinomycin was added to each well. The plate was shaken for 5 sec, and then fluorescence measurements were made every 20 sec for 2 min.

### Protein thermal shift assay

The protein thermal shift assay was performed following previously published procedures^40^. Briefly, recombinant proteins were diluted in PreScission Protease buffer with vector (1.25% [v/v] DMSO in 1x PBS), 40 μM of methotrexate (catalog #454126; MilliporeSigma), or 10 mM MnCl_2_ to reach a final protein concentration of 0.2 mg/mL. After 10 min of incubation on ice, SYPRO Orange Protein Gel Stain (catalog #S6650; Thermo Fisher Scientific) was added to reach a final concentration of 5x. 25 μL of the samples was loaded to each well on a low-profile non-skirted 96-well plate (catalog #AB-0700; Thermo Fisher Scientific), and a qPCR optical grade plate seal (catalog #Ab-1170; Thermo Fisher Scientific) was used to seal the plate. A CFX96 Touch Real-Time PCR Detection System (Bio-Rad) was used to measure the fluorescence signals from 10°C to 95°C, and the CFX Maestro software (Bio-Rad, version 2.3) was used for data acquisition following the protocol from Bio-Rad.

### Phosphatase activity assay

d32-PPTC7 (N-terminal truncation of the first 32 amino acids of H. sapiens PPTC7) was expressed in the pGEX-6P-1 vector and purified as described above. 4 μM of d32-PPTC7 was added incubated with 10 mM of the generic phosphatase substrate para-nitrophenyl phosphate (pNPP, New England Biolabs P0757S) and 10 mM MnCl_2_ diluted to a final volume of 100 μL of 50 mM TRIS, pH 8.0. The addition of MnCl_2_ to a final concentration of 10 mM was used to initiate the reaction. A reaction lacking MnCl_2_ served as a negative control. The dephosphorylation of pNPP into the chromogenic product of para-nitrophenol (pNP) was measured by monitoring the absorbance at 405 nm on an Epoch2 plate reader (BioTek) controlled by Gen5 software (version 3.10). Reactions were run at room temperature for 30 minutes, with the maximal slope of a minimum of 5 points used to calculate relative enzyme activity.

### Statistical analysis and software

Data processing was done in Excel (Microsoft), and statistical analysis was done in Prism (GraphPad, version 11.0.2). Ordinary one-way ANOVA was performed for Figure 4E, and ordinary two-way ANOVA was performed for Figure 4O, where data was assumed to follow a Gaussian distribution but not formally tested. All scatter plots and bar graphs were generated using Prism, then imported into Affinity Designer 2 (Serif, version 2.6.5) to make the finalized figures.

### Disordered protein analysis and design

Disorder was predicted using metapredict (V3)^81^. Helical potential was predicted with SPARROW using the dssp_helicity() predictor in probability mode. The helical potential predictor is a per-residue classification model, implemented as a PARROT-based^82^ long short-term memory bi-directional recurrent neural network (LSTM-BRNN). The model was trained to predict the binary DSSP helicity labels on a subset of structures from the AlphaFold2 database^83^. Version 0.2.2 of SPARROW was used for predictions and is available at https://github.com/idptools/sparrow with documentation at https://idptools-sparrow.readthedocs.io/en/latest/. Disordered variants were designed using GOOSE, which is available at https://github.com/idptools/goose with documentation at https://goose.readthedocs.io/en/latest/^52^. Code for IDR design is available at https://github.com/holehouse-lab/supportingdata/tree/master/2026/yan_and_wei_2026.

