## Supplemental Figures for "Combinatorial sequence elements fine-tune mitochondrial protein import to facilitate dual localization"

Yan and Wei et al.

**Supplemental Figure 1**

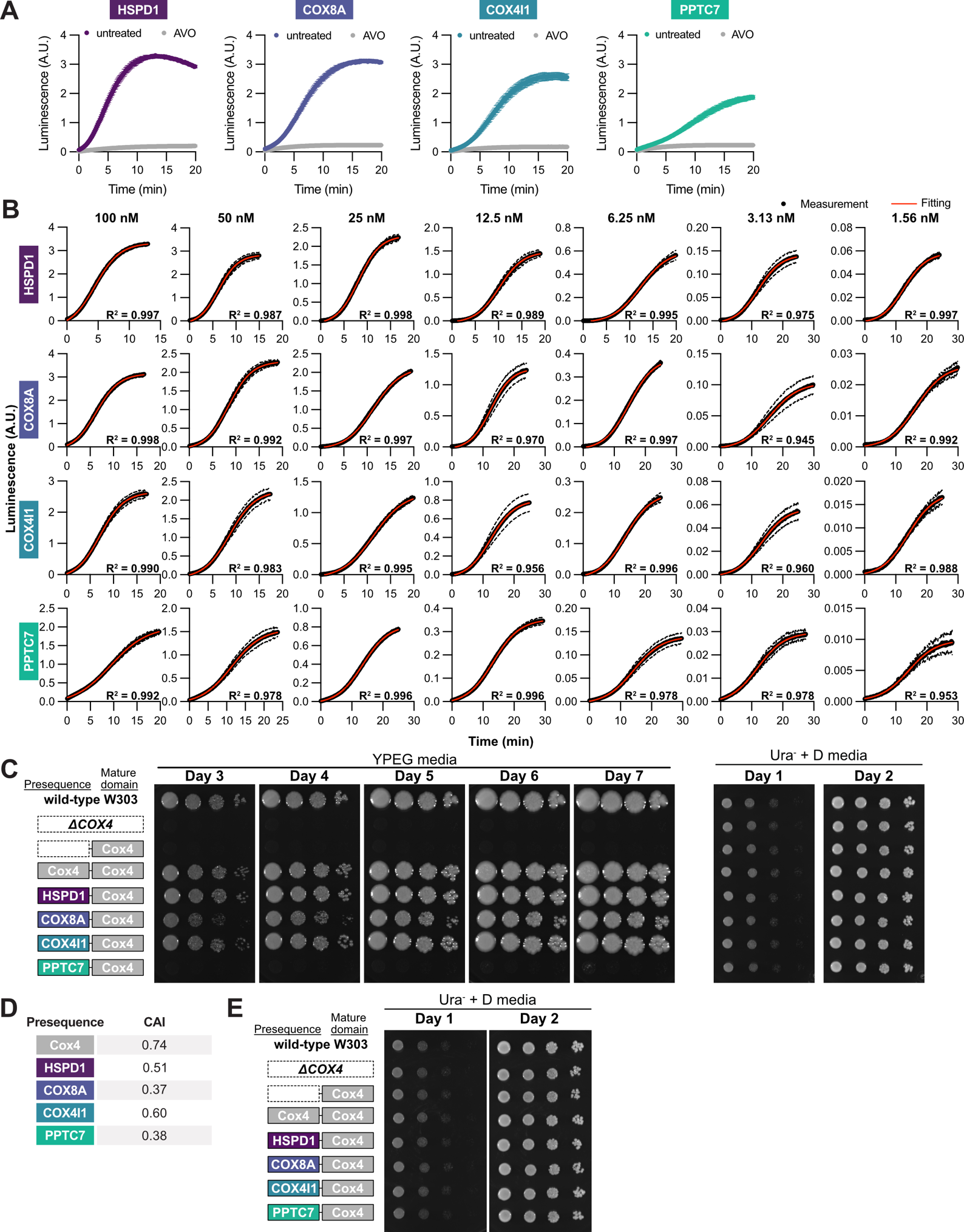

**Supplemental Figure 1**

**(A)** MitoLuc measurements about the import of the four presequence-DHFR-HiBiT fusion proteins with or without treating the mitochondria with antimycin A, valinomycin, and oligomycin. n = 3 independent measurements. Data represented as mean ± standard deviation.

**(B)** Import traces of the four presequence-DHFR-HiBiT fusion proteins from the MitoLuc assay were fitted with the general logistic function at each precursor concentration. n = 3 independent measurements. Data represented as mean ± standard deviation.

**(C)** Spotting assay that compared the growth of wild-type W303 yeast and *ΔCOX4* yeast rescued with vector or Cox4 mature domain fused to varied presequences (without codon optimization). Serial dilutions of yeast were plated on a YPEG (3% ethanol, 3% glycerol) plate or a Ura^-^ (2% glucose) plate and incubated at 30°C for the indicated number of days.

**(D)** Codon adaptation index of the genomic DNA sequences of the four human presequences and yeast Cox4 presequence calculated for expression in *S. cerevisiae*.

**(E)** Spotting assay comparing the growth of wild-type W303 yeast and *ΔCOX4* yeast rescued with vector or Cox4 mature domain fused to varied presequences (with codon optimization for COX8A and PPTC7). Serial dilutions of yeast were plated on a Ura^-^ (2% glucose) plate and incubated at 30°C for the indicated number of days.

**Supplemental Figure 2**

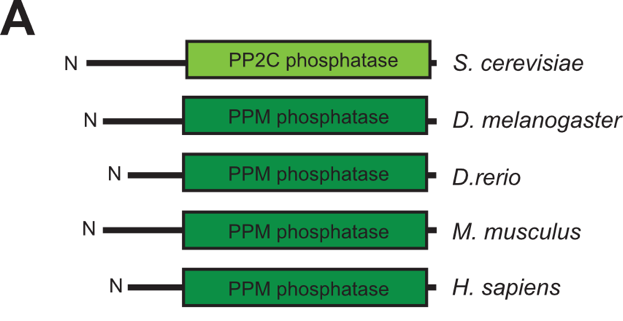

**Supplemental Figure 2**

Domain structure of PPTC7 orthologs across evolution.

**Supplemental Figure 3**

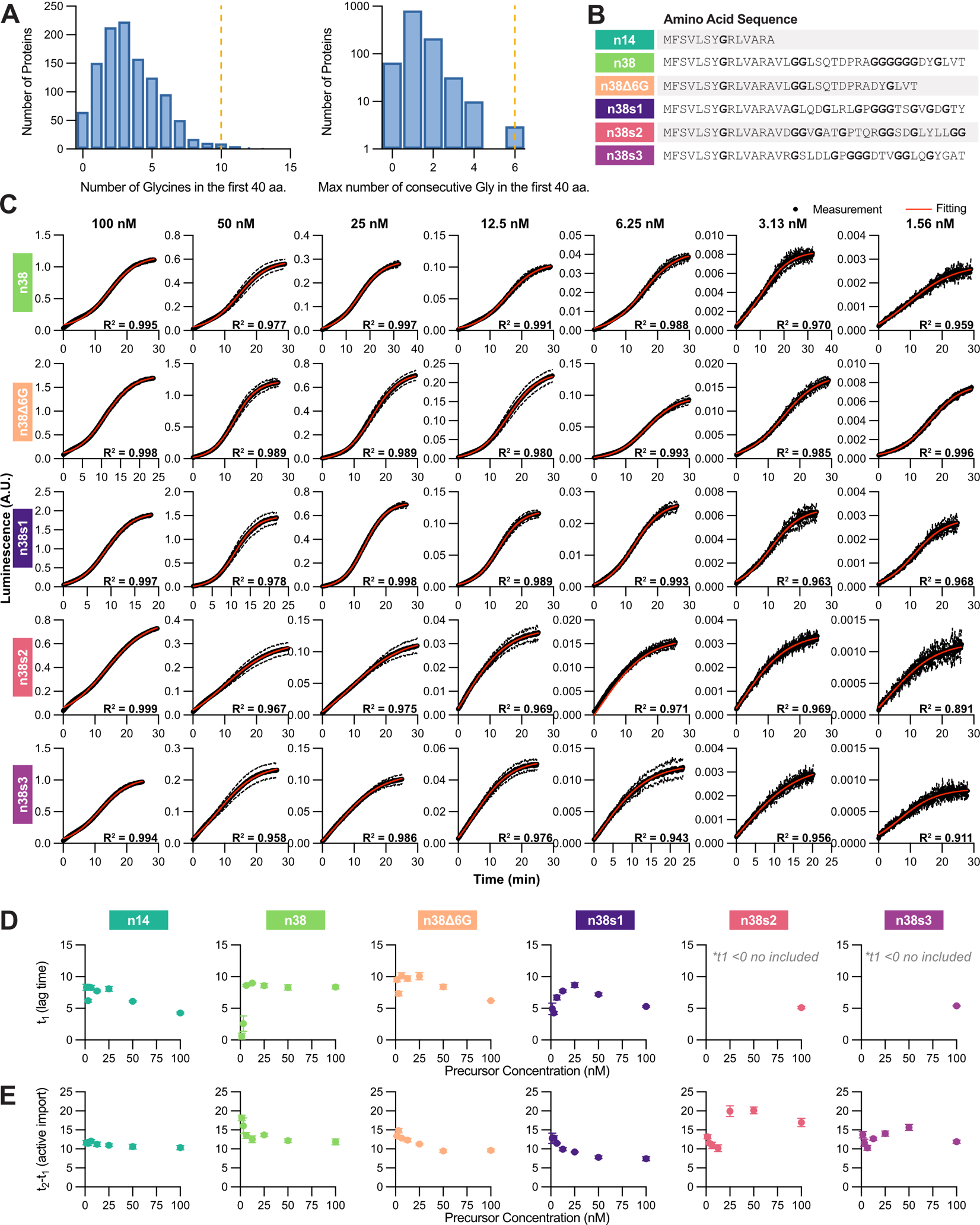

**Supplemental Figure 3**

**(A)** Distribution of mitochondrial proteins in terms of the number of glycines in the first 40 amino acids or the maximal number of consecutive glycines in the first 40 amino acids. The orange dotted line corresponds to the position of PPTC7.

**(B)** Amino acid sequences of the PPTC7-derived sequences tested in Figure 3.

**(C)** Import traces of the five PPTC7-DHFR-HiBiT fusion proteins from the MitoLuc assay were fitted with the general logistic function at each precursor concentration. n = 3 independent measurements. Data represented as mean ± standard deviation.

**(D-E)** Parameters extracted from the fitting shown in (C). Fitting results for lag time (D) and duration of active import (E) are shown as a function of precursor concentration. For the lag time, negative values are omitted. If t_1_ < 0, duration of active import equals t_2_ rather than t_2_-t_1_. n = 3 independent measurements. Data represented as mean ± standard deviation.

**Supplemental Figure 4**

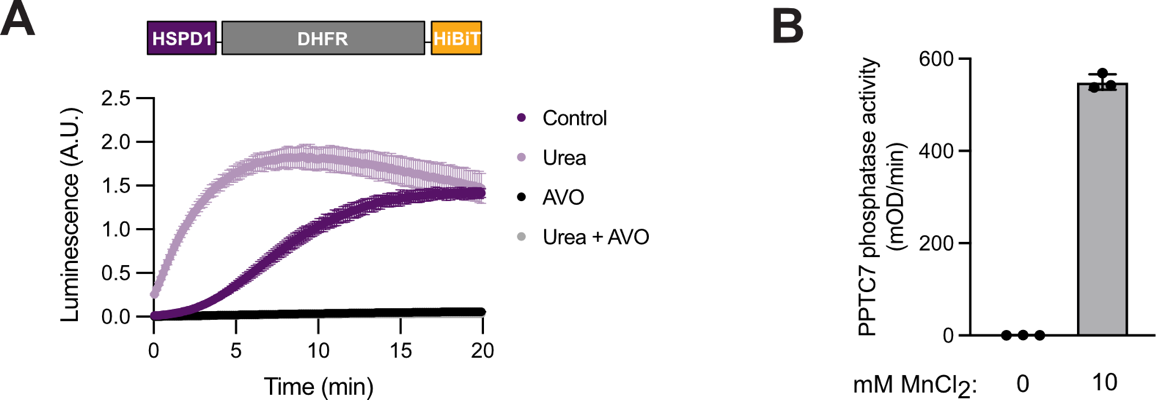

**Supplemental Figure 4**

**(A)** MitoLuc measurement of the import of 25 nM HSPD1-DHFR-HiBiT with and without urea denaturation into mitochondria with and without AVO treatment. n = 3 independent measurements. Data represented as mean ± standard deviation.

**(B)** Enzyme assay of recombinant 4 μM PPTC7Δ32 in the absence (0 mM) or presence (10 mM) of MnCl2. Phosphatase activity measured as rate of formation of para-nitrophenol (pNP) absorbance at 340 nm. . n = 3 independent measurements. Data represented as mean ± standard deviation.
